# Shifts in gut microbiome and metabolome are associated with risk of recurrent atrial fibrillation

**DOI:** 10.1101/2020.01.26.920587

**Authors:** Kun Zuo, Jing Li, Jing Zhang, Pan Wang, Jie Jiao, Zheng Liu, Xiandong Yin, Xiaoqing Liu, Kuibao Li, Xinchun Yang

## Abstract

Specific alterations of gut microbiota (GM) in atrial fibrillation (AF) patients, including elevated microbial diversity, particularly perturbed composition, imbalanced microbial function, and associated metabolic pattern modifications have been described in our previous report. The current work aimed to assess the association of GM composition with AF recurrence (RAF) after ablation, and to construct a GM-based predictive model for RAF. Gut microbial composition and metabolic profiles were assessed based on metagenomic sequencing and metabolomic analyses. Compared with non-AF controls (50 individuals), GM composition and metabolomic profile were significantly altered between patients with recurrent AF (17 individuals) and the non-RAF group (23 individuals). Notably, discriminative taxa between the non-RAF and RAF groups, including the families *Nitrosomonadaceae* and *Lentisphaeraceae*, the genera *Marinitoga* and *Rufibacter*, and the species *Faecalibacterium* sp*. CAG:82*, *Bacillus gobiensis*, and *Desulfobacterales bacterium PC51MH44*, were selected to construct a taxonomic scoring system based on LASSO analysis. An elevated area under curve (0.954) and positive net reclassification index (1.5601) for predicting RAF compared with traditional clinical scoring (AUC=0.6918) were obtained. The GM-based taxonomic scoring system theoretically improves the model performance. These data provide novel evidence that supports incorporating the GM factor into future recurrent risk stratification.

## Introduction

Atrial fibrillation (AF), the commonest arrhythmia among human cardiac diseases, is considered to cause a heavy global burden and directly impair the patient’s quality of life [1]. The morbidity rate of AF is becoming increasingly high worldwide, and the affected patients have approximately two-fold mortality rate increase compared with individuals with sinus rhythm due to cardiac and cerebrovascular events, such as cerebral stroke [2]. Medical management of AF with antiarrhythmic medications yields only partial effectiveness and is often associated with multiple adverse effects [3]. Hence, percutaneous radiofrequency catheter ablation represents an important treatment strategy and option for AF patients, especially individuals showing intolerance or symptomatic disease refractory to Class I or III antiarrhythmics [4]. Another issue is therefore raised, as ablation still carries the possibility and risk of AF recurrence (RAF). Not all AF patients that undergo radiofrequency catheter ablation remain in stable sinus rhythm. The success rates of catheter ablation maintaining sinus rhythm and avoiding a recurrence of AF after ablation are hard to predict and control [5–7]. To date, multiple variables, such as left atrial diameter and N-terminal pro–B-type natriuretic peptide (NT-proBNP), are considered risk factors for the recurrence of AF upon catheter ablation; however, these biomarkers lack specificity, and their predictive powers are barely satisfactory [8, 9]. The clinical scoring system, including CAAP-AF (coronary artery disease [CAD], age, left atrial size, persistent AF, unsuccessful antiarrhythmics, and female gender), DR-FLASH (diabetes mellitus, abnormal renal function, persistent type of AF, LA diameter > 45 mm, age > 65 years, female gender, and hypertension), and APPLE (age > 65 years, persistent AF, abnormal estimated glomerular filtration rate [eGFR; < 60 ml/min/1.73 m^2^], as well as LA diameter above 43 mm and ejection fraction below 50%) scores, could provide a realistic AF ablation outcome expectation for individual patients [7, 10–13]. However, this scoring system is simple and requires further modifications for increased robustness via substitution of etiologic factors by surrogate variables. Consequently, developing a novel and better predictive model, which would help physicians identify which individual patients could really benefit from catheter ablation and choose the best treatment option, is quite important.

The alteration and potential function of the gut microbiota in various pathologies, either as diagnostic biomarkers or contributors to pathogenesis, have attracted increasing attention [14, 15]. For example, the gut bacterium *Fusobacterium nucleatum* has been identified as a potent biomarker for improving the diagnostic performance of the fecal immunochemical test (FIT), helping detect tumors otherwise missed by FIT. This could allow a step forward in designing a non-invasive, potentially more accurate, and cost-effective diagnostic tool for advanced colorectal neoplasia. Elevated amounts of species of the phylum *Bacteroidetes* are associated with prevention of checkpoint-blockade-induced colitis. Therefore, the identification of gut microbiota (GM)-based markers might help develop approaches for reducing the risk of inflammatory complications upon administration of cancer immunotherapeutic drugs [16, 17]. Recently, we have assessed the role of GM in AF. Our team characterized the associations of GM alterations and metabolic patterns with AF by revealing the altered GM and bacteria-associated metabolites identified in AF cases in a previous research [18]. We further designed a random forest disease classifier based on abundances of co-abundance gene groups as variables for building a microbiota-dependent discrimination model for AF detection. The results showed that the differential gut microbiome signature could help diagnose AF, and the co-abundance gene groups originating from *Eubacterium*, *Prevotella*, *Ruminococcus*, *Dorea*, *Blautia*, *Bacteroides*, and *Lachnospiraceae* were essential in separating AF cases from control individuals. However, the correlation between altered GM and AF recurrence post-ablation remains unclear. Given the significance of GM shifts in AF patients as previously reported by our team, as well as the risk of AF recurrence following radiofrequency catheter ablation, we wondered whether the GM factor could be applied in predicting the risk of RAF, identifying patients who might benefit more from catheter ablation. Here, we evaluated the profiles of GM and metabolic patterns, assessed AF recurrence after radiofrequency ablation, and constructed a GM-dependent signature to identify the risk of AF recurrence.

## Results

### Characteristics of the study population and follow-up

In the current study, we included 90 participants from our previous cohort [18], with 50 non-AF controls (CTRs) and 40 AF patients. All AF cases underwent radiofrequency catheter ablation before feces collection, and the patients remained in sinus rhythm (normal beating of the heart) until the end of the procedure, with confirmed circumferential pulmonary vein isolation (CPVI) and ablation line blockage [4]. The occurrence of RAF was regarded as the endpoint. RAF was defined as any episode of non-sinus atrial tachyarrhythmia, as reported previously [19]. The patients with recurrent AF after the ablation would be allocated to the RAF group. And the patients without recurrence to the non-RAF group. To date, these AF patients have been followed-up for 15.6 ± 12.57 months. RAF was documented in 17 AF patients, with a postoperative recurrence rate of 42.5%. The clinical features of patients assessed in this study are summarized in Table 1. Briefly, compared with the non-AF CTR group, AF patients showed older age, higher incidence of type 2 diabetes mellitus (T2DM), reduced serum total cholesterol amounts, and higher intake frequency of drugs such as statins, dimethyl biguanide (DMBG), angiotensin receptor blockers (ARB), angiotensin-converting enzyme inhibitors (ACEI), and amiodarone. The baseline clinical characteristics of the non-RAF and RAF groups were comparable in terms of age, gender, body mass index (BMI), type 2 diabetes mellitus, hypertension and fasting blood glucose, serum creatinine, total cholesterol, and bilirubin amounts. In addition, we determined the CAAP-AF, DR-FLASH, and APPLE scores, which were significantly higher in the RAF group compared with the non-RAF group (Table 1; p=0.043, p=0.559 and p=0.564 for the CAAP-AF, DR-FLASH and APPLE scores, respectively). Furthermore, to assess the predictive values of the risk scores, receiver operating curve (ROC) analyses were performed. The results showed that the CAAP-AF score exhibited a higher area under curve (AUC) of 0.6981 than the DR-FLASH (AUC=0.5575) and APPLE (AUC = 0.4464) scores in predicting RAF. Therefore, the CAAP-AF score was selected to reflect clinical risk factors.

### AF recurrence is associated with the dynamically advanced degree of GM dysbiosis

The diversity index indicates the variety and richness of microbial entities in the gut, and is known to be associated with different disease states [20–22]. To assess gut microbial diversity in RAF patients, total gene amounts (Fig. 1a); alpha (within the individual) diversity, comprising Shannon’s index (Fig. 1b), Chao 1 richness (Fig. 1c), and Pielou’s evenness (Fig. 1d); and beta (between individuals) diversity of principal component analysis (PCA) (Fig. 1e), principal coordinate analysis (PCoA) (Fig. 1 f), and non-metric dimensional scaling (NMDS) (Fig. 1g) plots were analyzed at the species level based on metagenomic sequencing data. Compared with the non-AF CTR group, the non-RAF and RAF groups had significantly altered alpha and beta GM diversity, except for the Chao 1 index in the non-RAF group. Although no dynamic discrepancy in microbial diversity was observed between the non-RAF and RAF groups, there was an increasing and aggravated tendency of shifts in the RAF group, suggesting that patients experiencing recurrence after ablation might possess a more advanced degree of GM dysbiosis than non-RAF individuals.

**Figure 1.**
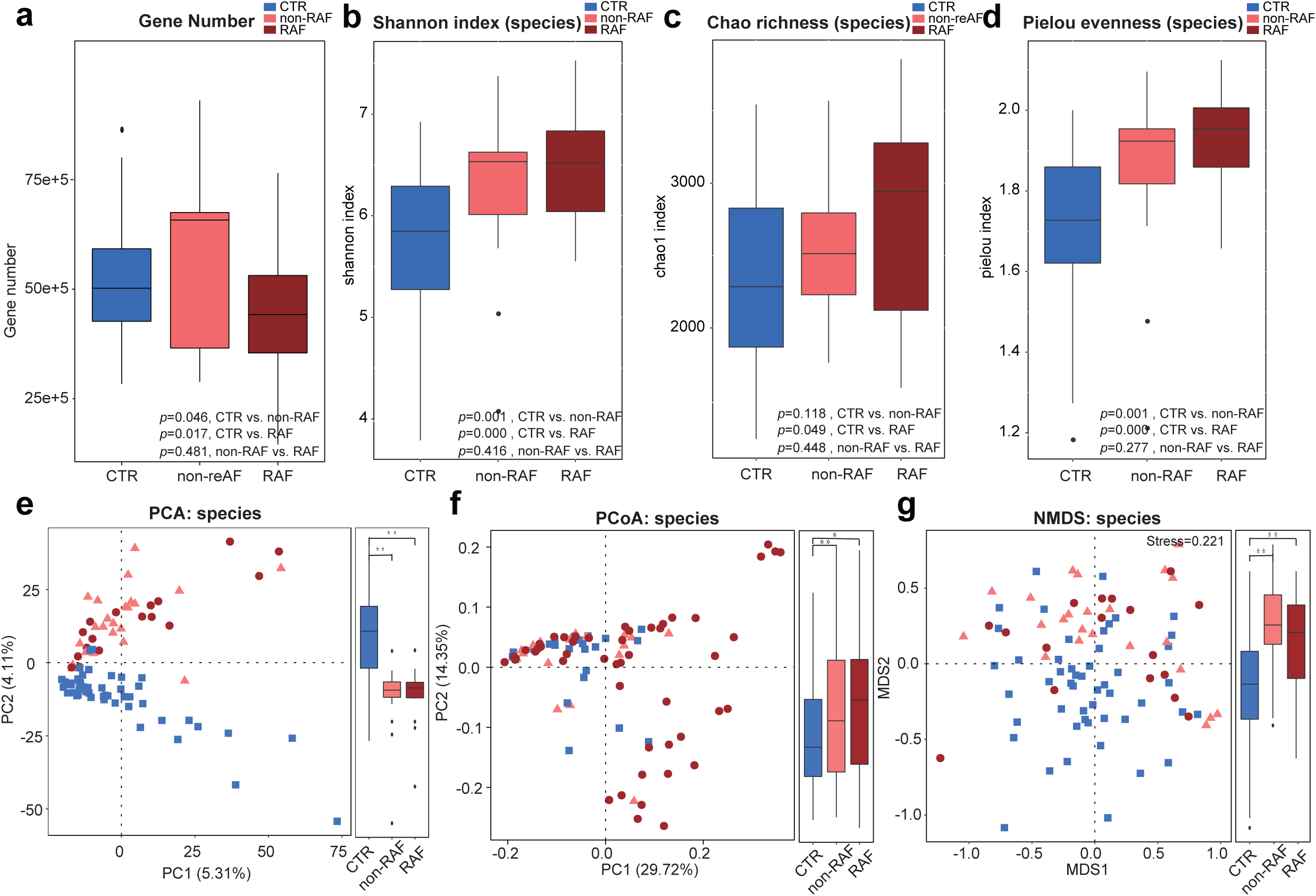
AF recurrence is associated with the dynamically advanced degree of dysbiosis in the GM. Gene number (a) and within individuals (alpha) diversity comprising Shannon index (b), Chao richness (c), and Pielou evenness (d) according to the species profile in non-AF CTR, non-RAF and RAF patients. Boxes are interquartile ranges, with lines denoting medians and circles being outliers. Between individuals (beta) diversity comprising PCA (e), PCoA (f), and NMDS (g) according to species abundances. The results depicted a dynamically increasing tendency of diversity among control, non-RAF and RAF cases. Blue squares represent non-AF CTR, pink triangles refer to non-RAF, and red circles denote RAF.

Meanwhile, considering the differences in clinical features such as age, T2DM incidence, total cholesterol amounts, and glutamic-pyruvic transaminase levels as well as medications administered between the AF and non-AF CTR groups, PCA was performed to examine whether the above GM profile changes in the non-RAF and RAF groups were actually driven by these discrepant clinical factors [23, 24]. The results in Fig. S1 showed that the dots representing individuals of different ages, T2DM diagnosis, total cholesterol, glutamic-pyruvic transaminase, or medication in the various plots were mixed together and failed to cluster into separate groups, indicating the low effects of these factors on the overall findings regarding the GM (Additional files 1: Fig. S1).

### Altered gut taxonomic profiles are associated with post-ablation RAF

The elevated gut microbial diversity and increased degree of GM dysbiosis in RAF indicate the possible overgrowth of some harmful microbes [25, 26]. Thus, the phylogenetic signatures of the GM were analyzed with the aim of further examining the changes in GM composition in RAF more specifically (Additional files 2-3: Table S1-2). Overall, the non-RAF and RAF groups shared most microbes detected in the non-AF CTR group, with 1219 genera (Additional files 3: Fig. S2a) and 5041 species (Additional files 3: Fig. S2d). Interestingly, some abundant bacteria, such as the genera *Faecalibacterium* and species *Faecalibacterium prausnitzii*, showed dynamically decreasing tendencies from non-RAF to RAF. In addition, *Ruminococcus* and *Eubacterium* exhibited progressively increasing trends from the non-RAF and RAF groups (Additional files 3: Fig. S2b, c, e, f). These progressive GM shifts associated with recurrent AF confirmed a dynamic and aggravating GM dysbiosis in patients who would suffer from recurrent AF after ablation.

Next, we assessed the taxa that were dramatically altered in the gut of non-RAF subjects and RAF patients at both the genus and species levels. Compared with the non-AF CTR group, a total of 354 and 337 genera, 1735 and 1646 species were significantly changed in the non-RAF and RAF groups, respectively (Additional files 5: Table S3). Generally speaking, the non-RAF and RAF groups shared 198 genera and 1077 species that were simultaneously altered (Fig. 2a, d), with most of these common bacteria exhibiting quite a similar tendency in the non-RAF and RAF groups (Fig. 2b, e). Several genera (e.g., *Prevotella*) and species (e.g., *Prevotella copri* and *Prevotella copri CAG:164*), which have been documented to be reduced in patients with Parkinson’s disease by a previous report [27], showed a decreased trend in the non-RAF group, and declined further in the RAF group. Consistently, genera such as *Ruminococcus*, *Blautia*, *Dorea*, and *Dialister*, as well as species including *Ruminococcus* sp. and *Dorea longicatena*, exhibited a progressively increased trend in patients experiencing recurrence post ablation (Fig. 2c, f). *Ruminococcus* exerts pro-inflammatory effects and contributes to inflammatory bowel disease (IBD) pathogenesis [28]. *Ruminococcus* transplantation in germ-free mice enhances interferon-γ, interleukin 17, and interleukin 22 amounts [29]. *Dialister* was shown to be associated with antepartum preeclampsia samples [30]. Thus, the balanced steady state in the gut is likely broken in AF subjects, especially in those who suffering from recurrence. The deficiency in health-promoting bacteria and the enrichment of disease-causing ones might be associated with RAF pathology after ablation.

**Figure 2.**
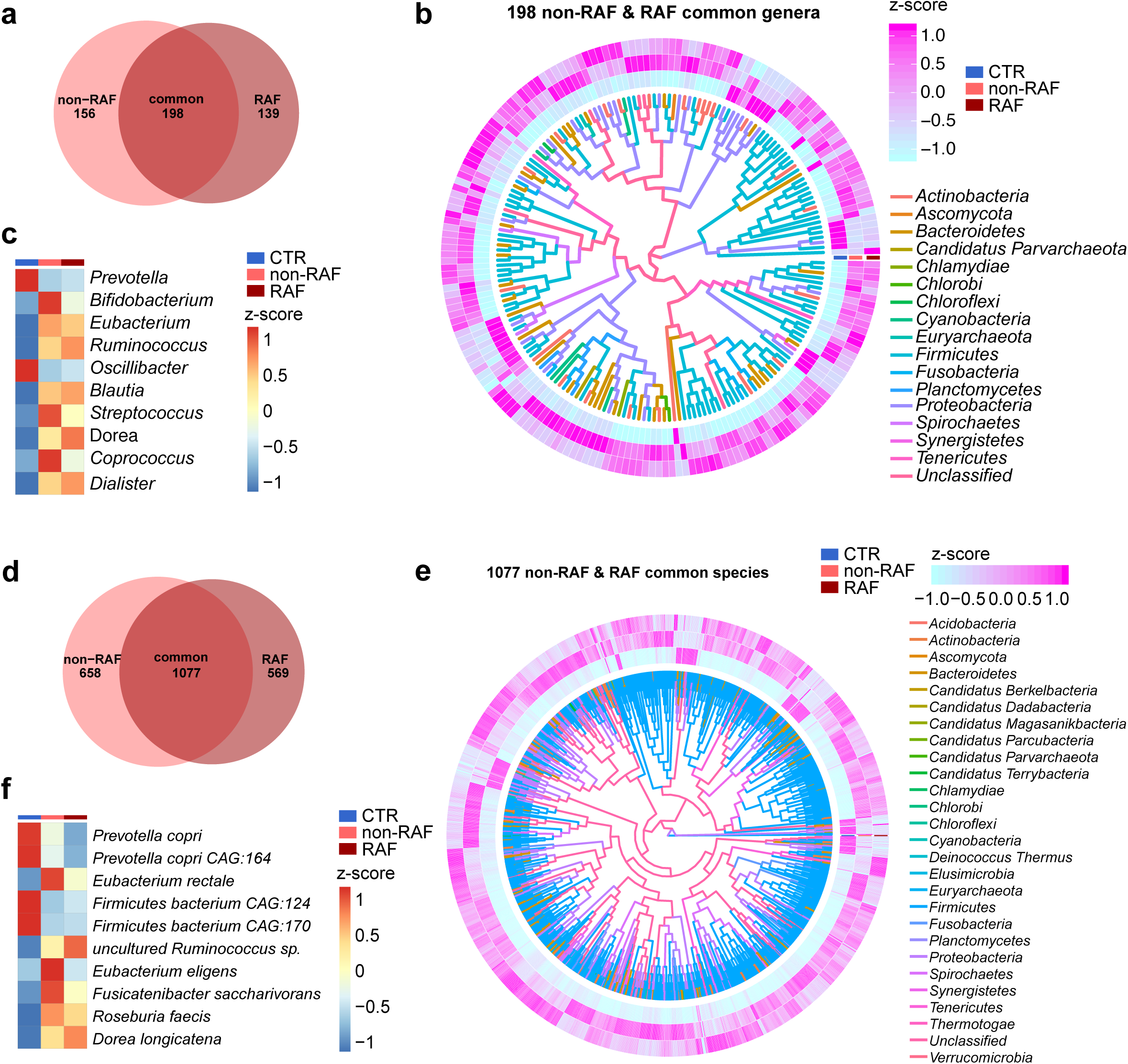
Common taxa in the non-RAF and RAF groups. a. Venn diagram showing the count of altered genera common to the non-recurrence of atrial fibrillation (AF) (non-RAF) (pink) and RAF (red) groups when compared to the non-AF control (CTR) group. The overlap revealed 198 genera simultaneously detected in AF patients with or without recurrence. b. Heat-map revealing 198 commonly altered genera in the non-RAF and RAF groups when compared to the non-AF CTR (*q* < 0.05 from Wilcoxon rank-sum test) and phylogenic associations. Abundance profile is reflected by the z-score, with genera grouped according to the Bray–Curtis distance. Negative (blue) and positive (pink) Z-scores reflect lower and higher abundance levels compared with the mean value, respectively. The colors of the lines inside denote the phyla of given genera. c. Heat-map of the first 10 shared genera (*q* < 0.05; Wilcoxon rank-sum test). Abundance profiles underwent transformation into Z-scores via average abundance subtraction and division by the standard deviation. Negative (blue) and positive (red) Z-scores reflected row abundance levels lower and higher compared with the mean, respectively. d. Venn diagram depicting the count of differential species common to the non-RAF (pink) and RAF (red) groups when compared with the non-AF CTR group. The overlap revealed 1077 species simultaneously detected in AF patients with or without recurrence. e. Heat-map depicting 1077 genera differentially present in the non-RAF and RAF groups when compared with non-AF CTR (*q* < 0.05 from Wilcoxon rank-sum test), and the corresponding phylogenic associations. Abundance profiles were plotted as z-scores, with genera grouped according to Bray–Curtis distance. Negative (blue) and positive (pink) Z-scores reflected row abundance levels lower and higher than the average, respectively. The colors of the lines inside denote the phyla of given genera. f. Heat-map of the first 10 shared species (*q* < 0.05; Wilcoxon rank-sum test). The abundance profiles were analyzed as in c. Negative (blue) and positive (red) Z-scores reflected row abundance levels lower and higher compared with the mean, respectively.

Besides the common shifts in gut taxa between the non-RAF and RAF groups, distinct alterations of bacteria profiles were identified exclusively in non-RAF or RAF patients. A total of 8 families, 4 genera, and 28 species showed significant differences between the non-RAF and RAF groups (Fig. 3). Bacteria such as *Methanobrevibacter* sp., *Methanobrevibacter smithii*, and *Methanobrevibacter curvatus* were more abundant in the RAF group, while microbes such as *Candidatus* sp., *Phycomycetaceae* sp., *Bacteroidetes bacterium RBG_13_43_22*, and *Faecalibacterium* sp. *CAG:82* were deficient in the non-RAF group. We speculated that the shared GM changes in the non-RAF and RAF groups might represent the core bacterial features of AF, and the unique shifts in GM composition in RAF patients might possibly account for the progression and recurrence of AF.

**Figure 3.**
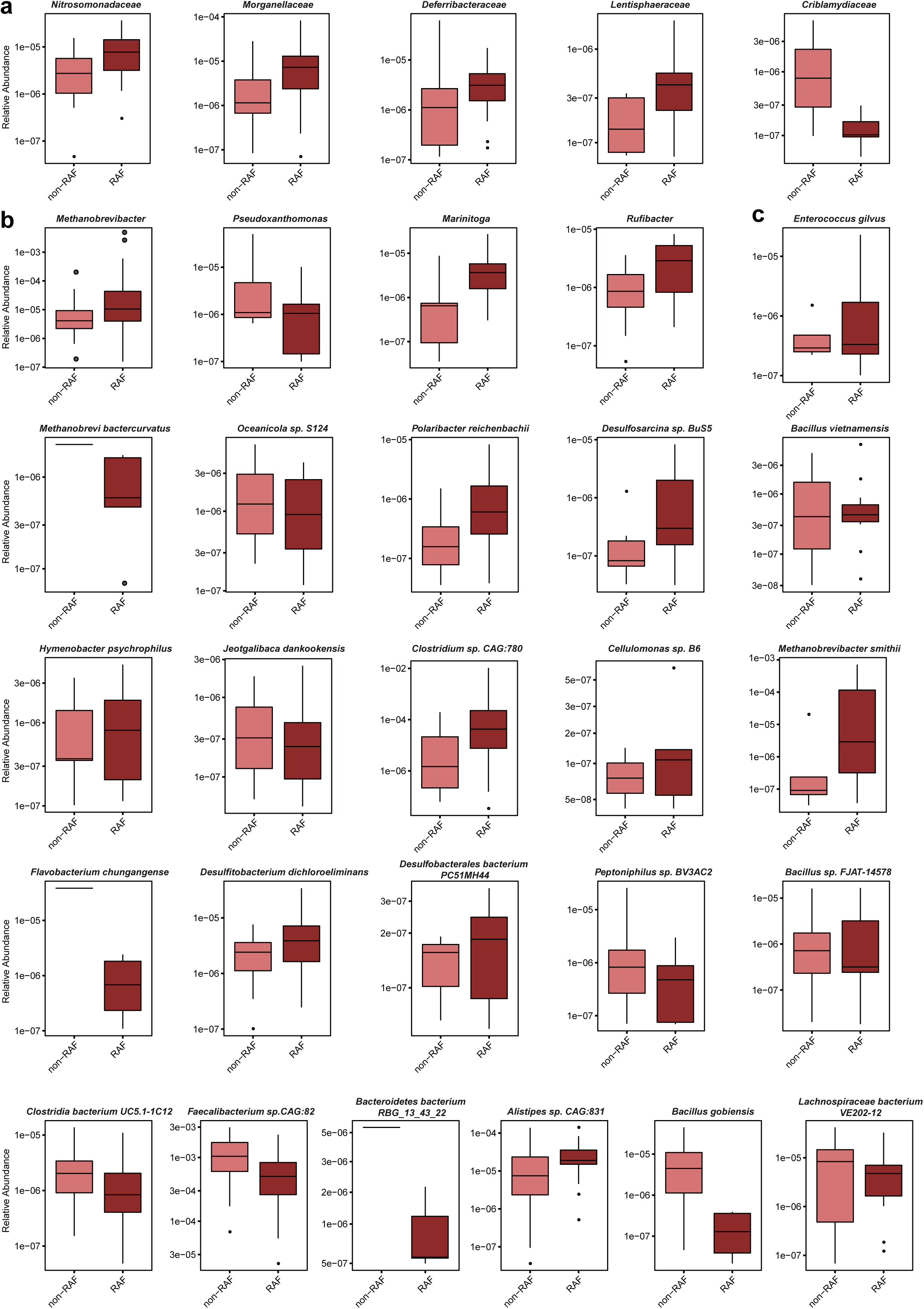
Distinctive taxa between non-RAF and RAF. Box plots of relative abundance (log transformed) of the distinctive taxa between individuals from non-RAF and RAF, including 5 families (a), 4 genera (b) and 22 species (c) at the criteria of q value <0.05; Wilcoxon rank sum test after filtering out in less than 10% of the samples detected and medians of zero. Boxes represent the interquartile ranges, lines inside the boxes mean medians and circles are outliers.

### GM functional profiles associated with RAF

To depict gut microbial gene functions among different AF categories, the Kyoto Encyclopedia of Genes and Genomes (KEGG) database was applied as described previously by our team [15, 18]. Briefly, the non-RAF and RAF groups could not be distinguished from each other, but were separated clearly when compared with non-AF CTRs by PCA, PCoA, and NMDS plots, which indicated drastic alterations in GM functions between the groups (Fig. 4a–c). Compared with non-AF CTRs, there were 201 KEGG modules in different enrichment shared between the non-RAF and RAF groups (adjusted p < 0.05, Wilcoxon rank-sum test, Fig. 4d; Additional files 6: Table S4). The majority of these functional modules also shared the same trends in the non-RAF and RAF groups (Fig. 4e). Some bacterial functions, including the citric acid cycle and amino acid biosynthesis, were deficient in the non-RAF and RAF groups, and are quite essential for human physiological health (Fig. 4e). Moreover, two KEGG modules, comprising the capsular polysaccharide transport system and xylitol transport system, were significantly elevated in the RAF group in compared with non-RAF individuals (Fig. 4f). However, the specific associations of these microbial functions with AF recurrence remain to be elucidated.

**Figure 4.**
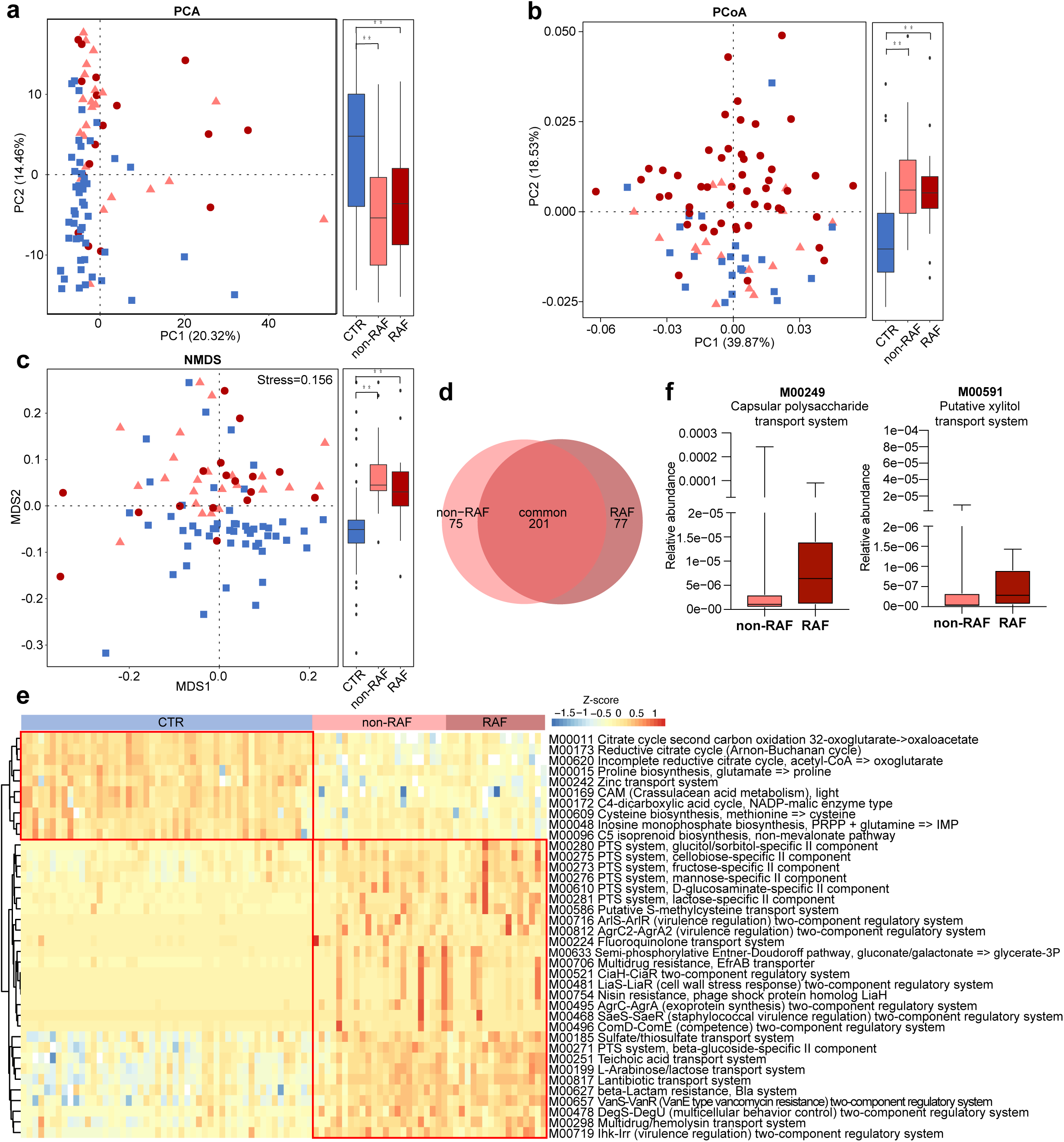
GM functional profiles associated with RAF. a, b, c. PCA (a), PCoA (b), NMDS (c) according abundance levels of KEGG modules showing disordered GM functional profiles in non-RAF and RAF cases. Blue squares represent non-AF CTR, pink triangles refer to non-RAF, and red circles denote RAF. d. Venn diagram depicting the count of differentially represented KEGG modules common in non-RAF (pink) and RAF (red) versus non-AF CTR. The overlap revealed 201 KEGG modules shared by non-RAF and RAF. e. Heat-map of 38 shared functional modules (q < 0.0001; Wilcoxon rank sum test). Abundance profiles underwent transformation into Z-scores via average abundance subtraction and division by the standard deviation. Negative (blue) and positive (red) Z-scores reflected row abundance levels lower and higher compared with the mean, respectively. f. Box plots of 2 differential KEGG modules between non-RAF (pink) and RAF (red) at the criteria of annotation in more than 20% of the samples. Box, interquartile range; line inside a box, median; circle, outlier.

### RAF is associated with disordered metabolomic profiles

The potential mechanisms mediating gut microbial function in human health rely on the interactions of gut microbe-derived metabolites with target organs [31, 32]. Therefore, metabolomic analyses based on LC-MS were performed to assess the metabolomic profiles of AF patients with or without the risk of AF recurrence following ablation. In this study, samples sufficient for metabonomic analysis were not obtained from all patients. Finally, a subset of 60 participants, comprising 36 non-AF CTRs, 13 non-RAF patients, and 11 RAF patients were included in serum metabolite analysis, and 52 individuals (17 non-AF CTRs, 20 non-RAF patients, and 15 RAF patients) were enrolled for metabolomic profile assessment in the feces. In serum, 2,500 and 1,733 features in the positive (ESI+) and negative (ESI−) ion modes were obtained, respectively. In fecal samples, 2,549 ESI+ and 1,894 ESI− features were observed. Overall, global metabolic changes in either serum or feces were revealed between the non-RAF and RAF groups by both partial least squares discriminant analysis (PLS-DA) and orthogonal PLS-DA (OPLS-DA) plots, with significant separation detected between non-RAF and RAF patients in modes of ES+ and ES− (Additional files 6: Fig. S3).

Overall, 94 circulating and 52 fecal metabolites showed simultaneous alterations in both non-RAF and RAF patients compared with non-AF CTRs (Fig. 5a). Interestingly, all 17 metabolites showing overlap in serum and stool specimens (Fig. 5 b, c, d) had similar variation trends in the non-RAF and RAF groups. Eight of them were synchronously altered in the serum and feces, and thus speculated to constitute the common features and core metabolites associated with AF development, which needs further investigation (Additional files 8: Table S5). In addition, two fecal metabolites, 7-methylguanine and palmitoleic acid, were found to be markedly reduced in cases with RAF in comparison with the non-RAF group. In addition, 7-methylguanine and palmitoleic acid showed no higher abundance in the CTR group compared with non-RAF and RAF cases (Fig. 5g).

**Figure 5.**
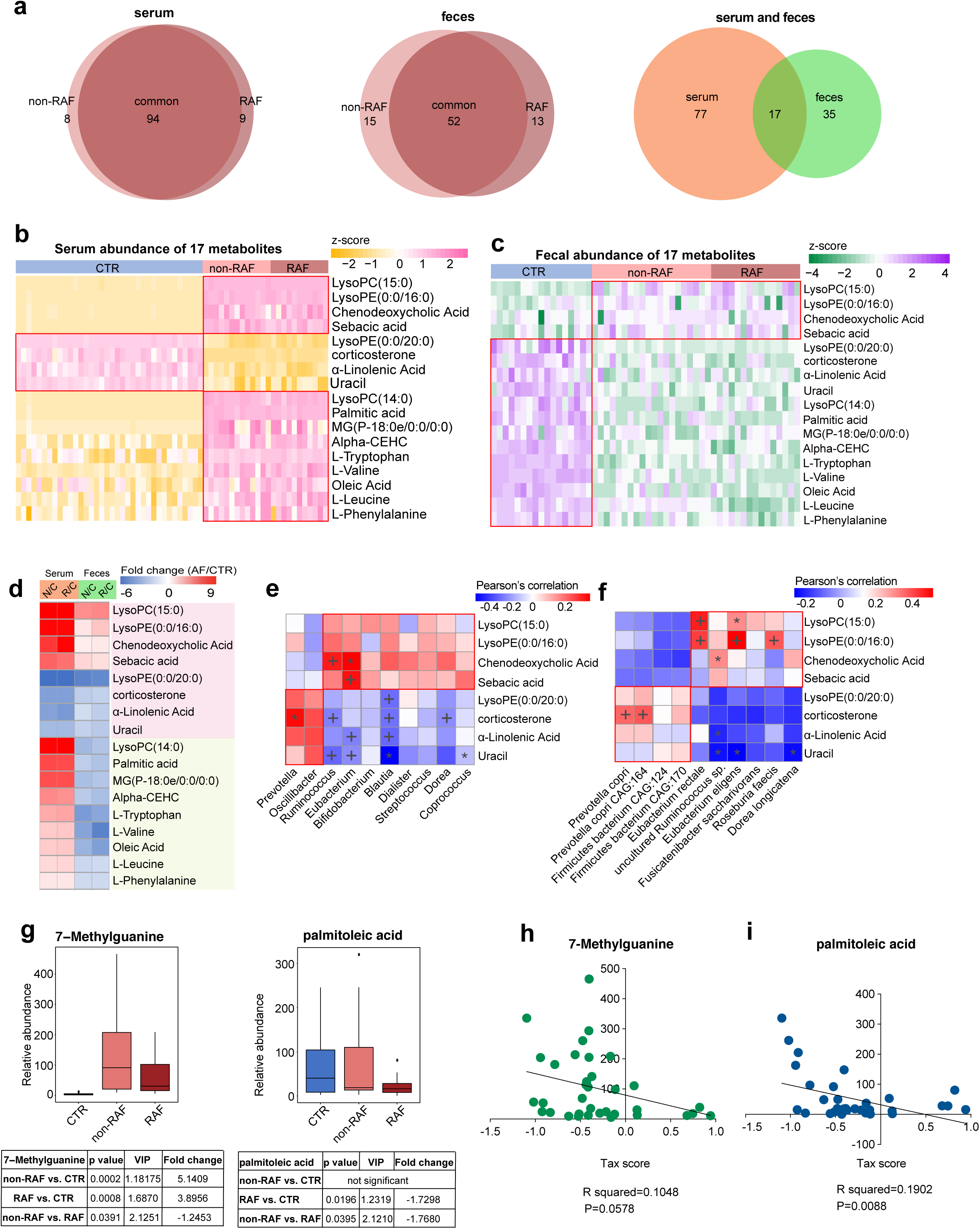
Abnormal metabolic patterns associated with recurrent AF. a. Venn diagram showing the amount of common differential metabolites in the non-RAF (pink) and RAF (red) groups when compared with the non-AF control (CTR). The overlap revealed 94 serum and 52 fecal metabolites simultaneously detected in the non-RAF and RAF groups, while 17 endogenous substances were simultaneously found in fecal and serum samples. b, c. Heat-map of 17 serum (b) and fecal (c) shared metabolites. Abundance profiles underwent transformation into Z-scores via average abundance subtraction and division by the standard deviation. Negative (yellow) and positive (pink) Z-scores reflected row abundance levels lower and higher compared with the mean, respectively. d. Heat-map depicting fold changes (AF/CTR) of 17 molecules with alterations in both serum and fecal specimens from AF cases. Fold changes underwent transformation into *t*-scores. Negative (blue) *t*-scores reflect compounds showing a decreasing trend in the non-RAF or RAF groups. Substances increasing or decreasing in both groups (n=8) or in a single group (n = 9) in fecal and serum specimens are depicted in pink and green, respectively. e, f. Relationship between eight simultaneously altered metabolites and the first 10 commonly detected genera (e) and species (f). Since the abundance levels of fecal metabolites mirrored those of GM-produced substances, fecal metabolomics data underwent Spearman’s correlation analysis. Blue, negative correlation; yellow, positive correlation, *p < 0.05, +p < 0.01. g. Box plots of two fecal distinctive metabolites between the non-RAF (pink) and RAF (red) groups. Box, interquartile range; line inside a box, median; circle, outlier. h, i. Correlation between taxonomic (Tax) score and two taxa distinctive between the non-RAF and RAF groups (R^2^=0.181, p=0.0023 for 7-methylguanine; R^2^=0.1217, p=0.014 for palmitoleic acid. Pearson linear correlations).

For assessing the associations of altered metabolites with changed GM, Pearson’s correlation analysis was carried out to evaluate the gut genera (Fig. 5e) and species (Fig. 5f) frequently changed in the non-RAF and RAF groups, in relation with the eight above-mentioned representative metabolites. Notably, the metabolites enriched in non-RAF and RAF patients, including lysophosphatidylethanolamine (LysoPE) (0:0/16:0), chenodeoxycholic acid (CDCA), and sebacic acid, were highly correlated with several AF-enriched genera (*Ruminococcus* and *Eubacterium*) and species, including *Eubacterium rectale*, *Roseburia inulinivorans*, and *Roseburia faecis*. Meanwhile, metabolites deficient in non-RAF and RAF patients, such as α-linolenic acid, were negatively correlated with AF-enriched genera, including *Eubacterium* and *Blautia*. Furthermore, correlation analyses between the two distinctive fecal metabolites and gut microbes showed that non-RAF enriched 7-methylguanine and palmitoleic acid were negatively associated with taxonomic (Tax) score decrease in non-RAF patients (Fig. 5h, i. 7-methylguanine: R^2^ = 0.1048, 95%CI: −0.593 to 0.01068, p = 0.0578; palmitoleic acid: R^2^ = 0.1902, 95%CI: −0.6717 to −0.1203, p = 0.0088, respectively). Based on the significant associations of metabolites with gut taxa, the possibility is raised that GM dysbiosis might cause a deficiency in select hear-protective metabolic products and/or producing deleterious compounds, either directly or indirectly, thereby contributing to the recurrence of AF post ablation. The GM could be, therefore, considered a latent risk factor for RAF.

### Development and validation of a predictive model based on GM signature and clinical scores for RAF

Subsequently, we sought to establish and assess a predictive model to help make individualized estimates of AF recurrence after ablation. Firstly, we selected the most predictive taxa for RAF by performing least absolute shrinkage and selection operator (LASSO) analyses [33, 34]. The results showed that seven bacterial strains consisting of two families (*Nitrosomonadaceae* and *Lentisphaeraceae*), two genera (*Marinitoga* and *Rufibacter*), and three species (*Faecalibacterium* sp*. CAG:82*, *Bacillus gobiensis*, and *Desulfobacterales bacterium PC51MH44*) among the candidate variables (taxon differing between the non-RAF and RAF groups) remained statistically significant, with nonzero coefficients based on 40 AF individuals (Fig. 6a, b). Then, we defined a risk score as the Tax score based on a linear combination of these seven taxa-based markers, and calculated the Tax score via weighting with their respective coefficients. The model was constructed as follows: Tax score = [-0.5104 × (Intercept)] + [35,896.6613 × *Nitrosomonadaceae*] + [564,576.2087 × *Lentisphaeraceae*] + [25.6052 × *Marinitoga*] + [71,729.3882 × *Rufibacter*] + [-236.5270 × *Faecalibacterium sp. CAG:82*] + [-6180.8888 × *Bacillus gobiensis*] + [730,762.9872 × *Desulfobacterales bacterium PC51MH44*] (Fig. 6c, d). Patients in the RAF group generally had significantly higher Tax scores (p = 3.5973e-08, Additional files 9: Table S6).

**Figure 6.**
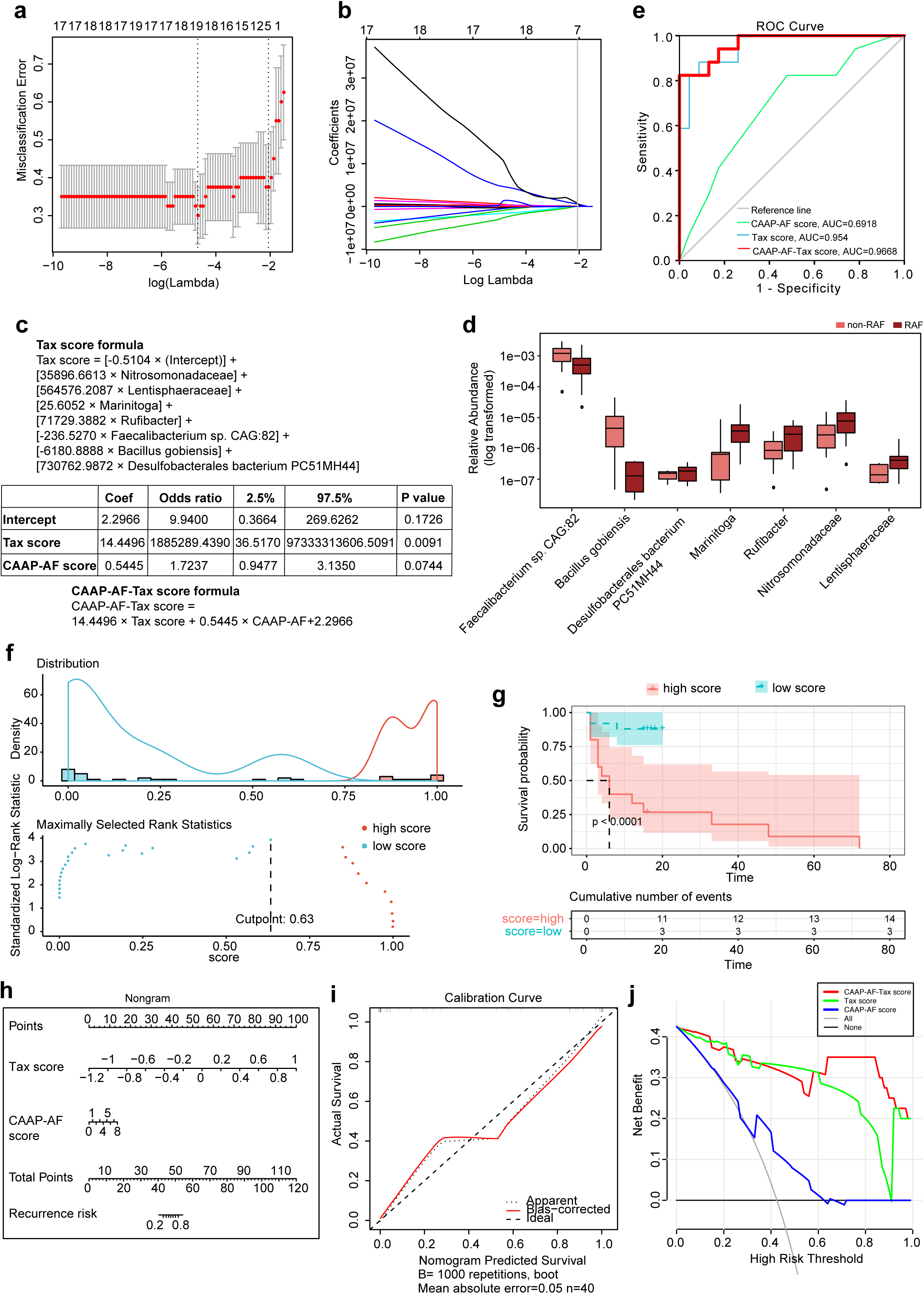
Taxonomic signature to predict recurrence following AF ablation. a. The tuning index (lamda) was selected in the LASSO model. Receiver operating characteristic curve generation was carried out, and its AUC was plotted against log (lamda). Dotted vertical lines depict the optimal values employing the minimum criteria and 1 standard error of the minimum criteria (-SE criteria). A lamda of 0.1267904, with log (lamda) of −0.8969136 was selected (1-SE criteria) based on the five-fold cross-validation method. b. LASSO coefficients of 37 taxonomic features. After excluding highly correlated (|r| ≥ 0.9) taxonomic features and linear combinations, 37 taxonomic features were retained. Coefficients were plotted versus log (lamda). A vertical line is shown at the value determined by five-fold cross-validation; optimal lamda yielded eight non-zero coefficients. c. The taxonomic (Tax) score was based on a linear combination of seven taxa-based markers, and calculated via weighting with their respective coefficients. Logistic regression analysis with the clinical CAAP-AF score and the developed Tax score was carried out using the enter method. A combined CAAP-AF-Tax score formula was constructed by weighting with the respective coefficients. d. Box plots of seven distinctive taxa between the non-RAF (pink) and RAF (red) groups. Box, interquartile range; line inside a box, median; circle, outlier. e. RAF is identifiable based on the Tax score or CAAP-AF score. Receiver operating curves for the CAAP-AF score, Tax score, and CAAP-AF-Tax score. The areas under the receiver operating curves (AUC values) were: CAAP-AF score, 0.6918 (95% confidence interval [CI]: 0.525–0.85, p=0.04); Tax score, 0.954 (95%CI: 0.8974–1.000, p=0.0055); CAAP-AF-Tax score, 0.9668 (95%CI: 0.9216–1.000, p=0.0011). f. Prognostic information provided by the CAAP-AF-Tax score model. Patients were ranked according to increased CAAP-AF-Tax score, and maximum difference in overall survival was obtained with a CAAP-AF-Tax score = 0.633286, splitting patients into high- and low-risk groups. g. Kaplan–Meier curves for overall survival prediction by the CAAP-AF-Tax score model. Cases were assigned to the high (red)- and low (green)-CAAP-AF-Tax score groups according to the corresponding cut-off CAAP-AF-Tax score value of 0.633286. There was a significant difference in overall survival between the high- and low-Tax score groups (p < 0.0001). h. Nonogram for recurrence risk prediction upon catheter ablation based on the Tax score. In the nomogram, each Tax score has a corresponding score on the score scale. A vertical line drawn down the score scale corresponding to the Tax score allows the risk of recurrence in a given patient to be easily and accurately read. i. Calibration curves of the Tax nomogram. Plots show calibrations for various models in terms of agreement between predicted and actual outcomes. Model performance is depicted by the apparent plot, and bias correction denotes the corrected value of the deviation, versus the 45-degree line representing the ideal prediction. j. Decision curve analysis of the Tax score-nomogram. The y-axis reflects the net benefit, with the red line representing the Tax score-nomogram; the grey and black lines denote the hypothetical cases with all and no cases exhibiting AF recurrence, respectively. At a threshold probability (patient or doctor) > 1%, employing the Tax score-nomogram for AF recurrence prediction shows elevated efficacy compared with the treat-all- or treat-none schemes. For instance, with an individualized threshold probability of 60% (a patient would be ineligible for therapy with a probability above 60%), a net benefit of 0.3125 is achieved in deciding whether to perform catheter ablation therapy.

In addition, to evaluate whether the gut microbiota (GM) signature could improve the predictive value of conventional risk factors such as clinical characteristics and drug usage, we performed univariate and multivariate Cox regression analyses, determining hazard ratios (HRs) and respective 95% confidence intervals (CIs) for parameters showing associations with AF recurrence upon ablation. The diagnostic performance of the model was evaluated by the C index [35]: 0.9–1.0, outstanding; 0.8–0.9, excellent; 0.7–0.8, acceptable [36].

Because of the limitation of sample size, the CAAP-AF score was selected as the synthetical reflection of numerous clinical characteristics (including age, gender, left atrial size, AF persistence, antiarrhythmics failed and CAD). We found that the Tax score and statin usage were significantly associated with RAF (Tax score, HR=2.5, 95%CI: 1.1-5.8, P=0.026; Statin usage, HR=4.8, 95%CI: 1.3-17, P=0.019). Meanwhile, other medication factors, including ACEI, ARB, CCB, β[blocker, propafenone and amiodarone administration were not significantly associated with RAF (Additional files 10: Table S7).

We next carried out multivariate-adjusted Cox regression based on the abovementioned indexes to assess whether GM could improve approaching utilizing conventional clinical factors. Thus, a clinical model incorporating the CAAP-AF score and statin usage, as well as a combined model including the CAAP-AF score, statin usage and Tax score were built (Table S8). After incorporating the clinical factors of RAF, Tax score retained a significant association with RAF incidence (HR=2.647, 95%CI: 1.038–6.749, P=0.041). Notably, the combined model had excellent (C index=0.8329, 95%CI: 0.7249-0.9410) and significant (p=0.0428) improvement in performance, in comparison with the clinical model (C index=0.7261, 95%CI: 0.5813-0.8709).

Then, using the enter method, logistic regression analysis with the clinical CAAP-AF score and the developed Tax score was carried out. The Tax score was identified as an independent predictor (Tax score: Coef=14.4496, Odds ratio [OR]=1,885,289.4390, 95%CI: 36.5170-97,333,313,606.5091; p=0.0091; CAAP-AF score, Coef=0.5445, OR=1.7237; 95%CI: 0.9477-3.1350; p=0.0744) (Fig. 6 c). Then, a combined predictive model containing two predictive scores, named CAAP-AF-Tax score, was constructed as follows: CAAP-AF-Tax score = 14.4496 × Tax score + 0.5445 × CAAP-AF + 2.2966, with values ranging from −12.4340 to 18.0986 (Additional files 10: Table S6). Patients with a score of −12.4340 and 18.0986 had predicted recurrence risks of 3.98e-04% and 99.9999986%, respectively (Fig. 6c).

To assess the predictive value of the CAAP-AF-Tax model, the AUC based on the ROC curve was determined and compared with those of the CAAP-AF and Tax scores. Notably, compared with the AUC for the CAAP-AF score alone (AUC=0.6918, 95%CI: 0.525-0.85, p=0.04), AUC for either the Tax score model (AUC=0.954, 95%CI: 0.8974-1.000, p=0.0055) or CAAP-AF-Tax model (AUC=0.9668, 95%CI: 0.9216-1.000, p=0.0011) was significantly higher (Fig. 6e). The predictive model was subsequently validated using the net reclassification index (NRI). The NRI after adding the CAAP-AF score to the Tax score was 1.1509 (p=0.0003), while that after adding the Tax score to the CAAP-AF score was 1.5601 (p=1.0735e-06). Therefore, the Tax score theoretically improved the CAAP-AF model for performance.

Then, the Kaplan–Meier method and log-rank test were carried out for assessing the prognostic capacities of the developed CAAP-AF-Tax model, and AF cases were assigned to high- and low-CAAP-AF-Tax score groups according to the optimal cut-off of 0.633286. There was a significant difference in overall survival between the high- and low-score groups (p < 0.0001, Fig. 6f, g).

To provide a quantitative method for predicting individual risk and RAF probability, a nomogram of the CAAP-AF-Tax model was established (Fig. 6h) and submitted to internal validation as previously reported [37]. The results showed C-index=0.9668 (95%CI: 0.9155–1). The calibration curve in Fig. 6i demonstrated good agreement with the probability of RAF. The Hosmer–Lemeshow test revealed non-statistical significance (p=0.6318) and suggested inconsiderable departure from the perfect fit [38]. Subsequently, in Fig. 6j, to determine the clinical value of the above Tax nomogram, we carried out decision curve analysis (DCA) via net benefit quantitation at various threshold probabilities [39]. We found that at a threshold probability (patient or doctor) of 1%, Tax score or CAAP-AF-Tax score nomogram use to predict RAF would provide more benefits compared with the treat-all- or treat-none schemes. Thus, the developed and validated predictive model might be a reliable method for RAF prediction, and help clinicians identify candidates who may benefit from future ablation therapy.

## Discussion

In the current study, we have acquired a series of intriguing results describing the profiles of altered GM and metabolic patterns in AF patients more likely to experience recurrence following radiofrequency ablation. The predictive model based on the gut taxonomic signature was built for identifying patients at high risk of RAF. We identified a gradually increasing degree of gut dysbiosis from non-AF to RAF, and found multiple bacteria simultaneously enriched in non-RAF and RAF patients. Meanwhile, imbalanced GM functions and metabolic alterations were observed, which indicates the possible GM function in eliciting AF recurrence via interactions with metabolites. This feature of GM is therefore suggested to be a potent risk biomarker for the selection of patients who would benefit from radiofrequency catheter ablation. It is recommended to include an additional focus on GM profiling in the future development of ablation risk stratification and strategies.

Strikingly, this study demonstrated that disordered GM constituted an independent risk factor for AF recurrence. Catheter ablation is an efficient therapeutic option for AF, and has been widely used in clinical settings. CPVI of extra–pulmonary vein (PV) AF substrate is the main strategy of ablation, which aims to suppress the arrhythmogenic substrate constituting the ablation target [4]. However, a high post-ablation recurrence rate calls for identifying novel markers that would enable an improved selection of patients who could benefit from ablation. PV reconnection and unrecognized or progressive extra–PV AF substrate constitute the immediate reflection of recurrence, but is neither necessary nor sufficient for RAF occurrence [40]. Therefore, based on the electro-physiological substrate behind AF persistence and progression, it remains difficult to identify beyond clinical AF parameters.

In our previous research, disordered GM was shown to be associated with the development of AF [18]. Interestingly, the present study indicated an incremental prognostic accuracy over clinical predictors of the novel predictive model based on GM taxonomic profiles. The newly defined Tax-score based on taxonomic profiles in the current work independently predicted AF recurrence, and findings from nomogram and decision curve analyses further confirmed its clinical value. Therefore, GM should be considered a potent predictive model for selecting patients for ablation, and additional focus on disordered GM profiles is strongly recommended in future ablation risk stratification. Although the number of samples included in the current research was small—and hence the robustness of the novel predictive model may not be strong enough—it provides preliminary results and offers a novel concept. Further validation studies with increased sample sizes could improve the overall universality.

Besides the distinctive taxa contained in the predictive score, two distinctive metabolites identified between non-RAF and RAF were significantly correlated with the Tax score. Although studies describing the direct protective effects of 7-methylguanine and palmitoleic acid against AF are scarce, investigators have identified the potential roles of 7-methylguanine and palmitoleic acid in the pathophysiological process related to AF. 7-methylguanine, a nucleotide contributing to the metabolic pathways of guanine-containing purines, is linked to cognition phenotype [41]. A recent study showed that 7-methylguanine was increased in incident type 2 diabetes millitus [42]. Meanwhile, palmitoleic acid has been suggested to enhance insulin sensitivity, stimulate insulin secretion, increase liver oxidation of fatty acids, improve blood lipid profile, alter the differentiation of macrophages [43], and improves metabolic functions in fatty liver tissue through peroxisome proliferator-activated receptor-α (PPARα)-dependent 5′ AMP-activated protein kinase (AMPK) activation [44]. Emerging evidence suggests that metabolic impairment is important for AF pathophysiology [45]. Notably, PPARα-dependent AMPK activation could result in suppressed inflammation [46]. Therefore, we speculate that reduction of fecal metabolites such as palmitoleic acid in RAF patients might contribute to excessive inflammation and facilitate AF recurrence. These GM-related metabolic changes may contribute to the progress of atrial tissue’s arrhythmogenic substrate aggravation following catheter ablation.

Given the associations of 7-methylguanine and palmitoleic acid with GM identified in the current work, these two metabolites are speculated to be potential players mediating the impact of GM dysbiosis on RAF progression, at least in part. Although it has not been assessed whether these two metabolites are directly produced by the GM, evidence demonstrating a link between 7-methylguanine, palmitoleic acid and GM is increasing. Palmitoleic acid belongs to the class of organic compounds known as long-chain fatty acids. Recent findings suggest that the metabolic activities of enteric microbiota may affect the levels of long-chain fatty acids [47]. Oral gavage of *Enterococcus faecalis* has been reported to affect a variety of long-chain fatty acids, including palmitoleic [48]. In addition, increased palmitoleic acid was previously observed in mice fed gut *Bifidobacterium breve* [49] and *Lactobacillus rhamnosus LA68* [50]. Moreover, palmitoleic acid is considered a metabolic phenotype biomarker in acute anterior uveitis patients due to its positive correlation with gut *Roseburia* [51]. 7-methylguanine belongs to the category of DNA adduct inducers [52]. The level of DNA adducts including 7-methylguanine and O^6^-methylguanine in the rat colon following administration of normal gut bacterial organisms has been previously investigated [52]; a slight reduction in the half-life of 7-methylguanine was observed after administration of *Lactobacillus acidophilus*. Based on the associations of fecal 7-methylguanine, palmitoleic acid and Tax score, the possibility was raised that GM dysbiosis might lead to deficiency in 7-methylguanine and palmitoleic acid in RAF patients, either directly or indirectly.

Inflammation is associated with multiple pathological events, including oxidative stress, apoptosis and fibrosis, which induce AF substrate generation. Therefore, low-grade inflammation is considered a potential mechanism contributing to AF. We found the species *Faecalibacterium sp. CAG:82* exhibited a decreased trend in the RAF group compared with non-RAFs. A previous study has demonstrated an anti-inflammatory effect of gut *Faecalibacterium* through inhibition of interleukin-6 and transcription 3/interleukin-17 pathway activation [53]. Therefore, it is speculated that reduced *Faecalibacterium* abundance in the intestine might increase various inflammatory cytokines, elicit low-grade inflammation, and thus lead to RAF. These interconnected microbial and metabolic changes suggest the involved microbes might contribute to AF recurrence through interactions with specific metabolites in the host.

In addition to the disparity between the non-RAF and RAF groups, similarities shared by these groups were also revealed in the current study, which may be more important and constitute key events in the onset—but not development—of AF. Furthermore, bacterial organisms and metabolites commonly altered in the non-RAF and RAF groups were significantly associated. CDCA and sebacic acid were found to be significantly correlated with several taxa. Elevated serum CDCA in the metabolic patterns of non-RAF and RAF patients has been indicated to have a critical function in the progress of structural remodeling in AF. CDCA is positively correlated with the left atrial low voltage area (LVA) and promotes apoptosis in atrial myocytes [54]. Furthermore, sebacic acid belonging to medium-chain fatty acids is significantly less abundant in both ulcerative colitis or Crohn disease patients [55]. Therefore, the shared GM and metabolic profile demonstrated above may be associated with or even contribute to AF onset.

These findings provided opportunities to take advantage of the GM for clinical application for improving GM-related AF pathogenesis [56, 57]. For example, utilizing fecal markers for identifying patients at high risk of RAF. Modulating microorganisms using antibiotics to inhibit disease-enriched bacteria [58], supplementing commensals [59, 60] or performing fecal microbiota transplantation to replenish disease-decreased bacteria are also recommended [61, 62]. Meanwhile, with the host–microbe crosstalk being in part mediated by bacterial metabolites, chemical approaches could be regarded as a promising therapeutic strategy. Dietary interventions, food compounds that can modify the GM (prebiotics) [59, 63] or metabolites generated from gut bacteria (postbiotics) [64, 65] might be beneficial. Engineering approaches like bacteriophages [66–68] specifically modifying gut bacteria are also suggested. These extensive findings will pave the way to translate GM use for clinical intervention, and more studies are imperative to evaluate its clinical value in the context of AF.

In conclusion, the present findings provide a comprehensive description of disordered GM profiles as well as its possible functions in AF patients with a high risk of recurrence following radiofrequency ablation. The importance and clinical value of bacterial taxonomic markers in patient selection for ablation are highlighted. More attention should be paid to disordered GM profiles while developing future ablation risk stratification strategies.

## Methods

### Study cohort

Forty non-valvular persistent AF (psAF) patients who underwent radiofrequency catheter ablation and 50 non-AF CTRs were included from our previous study [18]. Fecal samples were collected before radiofrequency ablation; the gut microbiome obtained before the ablation procedures was therefore utilized for predicting AF recurrence risk. Metagenomic sequencing results of 50 non-AF controls’ fecal specimens previously assessed by our team were employed as controls. For the 50 non-AF controls, exclusion criteria were: previous heart failure; CAD; structural heart disease; concurrent pathologies such as IBD, autoimmune disorders, liver disease, kidney diseases or malignancy; antibiotic or probiotic use within a month prior to enrolment. Patient baseline features were collected by face-to-face interviews and from hospital records.

### Catheter ablation and follow-up

In general, indications for AF ablation are symptomatic AF refractory or intolerant to one or more Class I or III antiarrhythmics, as well as symptom-free AF before administration of Class I or III antiarrhythmics [69]. A decision to perform ablation is taken upon careful consideration of complication risks, success rate, substitute options and patient preference.

Upon double-transseptal puncture under guidance of a 3D-electroanatomic mapping system (CARTO 3; Biosense Webster, Inc., Diamond Bar, CA, USA), a 3D-reconstructed image of left atrium was generated with a circular mapping catheter (NaviStarThermocool, Biosense Webster, Inc., Diamond Bar, CA, USA) followed by merging to 3D VR cardiac CT scan. Following circumferential pulmonary vein isolation (CPVI), linear ablation and complex fractionated atrial electrogram ablations were appended [4]. All catheter ablations were carried out by a single surgical team.

Following catheter ablation, patients underwent systematic follow-up and 12-lead electrocardiography at 3, 6, 12, and 18 months; respectively; an electrocardiogram would be recorded in case a patient complained of discomfort. Holter monitoring was performed at three- and six-months post-ablation and at six-month intervals afterwards. Antiarrhythmic medications were discontinued in the third month. AF recurrence was defined as any episode of non-sinus atrial tachyarrhythmia (atrial tachycardia, atrial flutter, or AF) lasting more than 30 s and occurring after the three-month post ablation blanking period [19]. The patients with recurrent AF after ablation would be allocated to the RAF group. And those without recurrence would be classified as the nonrecurrence (non-RAF) group.

In our electrophysiological team, antiarrhythmic medications would be discontinued at five half lives or more prior to ablation. After ablation, except for cases with contraindications (e.g., sinus bradycardia, second degree atrial-ventricular block, systolic blood pressure < 100 mmHg, hepatic dysfunction and dysthyroidism), all cases with persistent AF and some with paroxysmal AF would receive oral antiarrhythmic drugs for a 3-month period. In the current work, amiodarone and propafenone were administered to 30 (75%) and 7 (17.5%) individuals, respectively. Antiarrhythmic drugs were stopped in case of RAF recurrence following discontinuation for a period of time. During follow up, the patients who experienced RAF would resume with anti-arrhythmic medication; in case of persistent recurrent episodes in spite of drug administration, a second ablative surgery would be offered.

Furthermore, according to current guidelines [1], the risk of stroke in AF patients should be assessed by the CHA2DS2-VASc score. Generally, cases with no clinically proven risk factors for stroke do not require antithrombotic treatment. Meanwhile, those showing stroke’s risk factors (CHA2DS2-VASc scores ≥1 and ≥2 in males and females, respectively) without contraindications are recommended for oral anticoagulant (OAC) drugs. In our center, patients would take warfarin, dabigatran or rivaroxaban for ≥ 4 weeks before ablation. We carried out transesophageal echocardiography within 3 days prior to ablation. During the ablative surgery, heparin was administered before or right after transseptal puncture, adjusting the dose for achieving and maintaining an ACT ≥300 seconds. OACs should be administered for ≥ 3 months post-ablation. The continuation of OACs for more than 3 months post-ablation depends on stroke’s risk profile.

### GM assessment by Metagenomics

Whole-metagenome sequencing data of 90 fecal specimens assessed in the current study were obtained from a previous report by our team [18]. Bacterial DNA extraction utilized a TIANGEN kit (TIANGEN BIOTECH, China). Then, paired-end sequencing was carried out on an Illumina Novaseq 6000 (Illumina, USA), by Novogene Bioinformatics Technology (China), with an insert size of 300 bp and a read length of 150 bp. Metagenomic analyses followed the procedures described by our group [14, 15, 18]. Genes were predicted from various contigs with MetaGeneMark v12 (GeneMark, USA). Gene abundance was determined by numbering reads after normalization to gene length. Then, DIAMOND v0.7.9.58 was utilized for taxonomic assignments (default settings with the exception of −k 50 −sensitive −e 0.00001). Matches showing statistical significance for various genes with the first hit e≤10 × e-value were utilized for distinguishing taxonomic groups. The taxonomic levels of different genes were assessed with the MEGAN software (MEtaGenomeANalyzer); the abundance levels of different taxonomic groups were evaluated by summing up those of all included genes. The KEGG database (Release 73.1) was employed for gene alignment with DIAMOND (as described above) to evaluate the function of gut microbe. Proteins were assigned to the KEGG based on respective highest-scoring hits with ≥1 high-scoring segment pair comprising more than 60 hits. The abundance levels of KEGG modules were determined by summing up those of all genes included, respectively.

### GM assessment by metabolomics

Of the above 90 cases, metabolomic data of 60 serum and 52 fecal specimens were available [18]. Liquid chromatography-mass spectrometry (LC/MS) was carried out with a Hypercarb C18 column (Thermo Fisher, USA; 3 μm internal diameter, 4.6 mm×100 mm) on an UltiMate 3000 chromatography system (Thermo Fisher). Acetonitrile (Merck, Germany), methanol (Merck), formic acid (CNW, China), and DL-o-Chlorophenylalanine (GL Biochem, China) were used during the process. Data analysis was carried out as described in our previous reports [14, 18]. Partial least-squares discriminant analysis (PLS-DA) as well as orthogonal partial least-squares discriminant analysis (OPLS-DA) utilized the SIMCA-P software for clustering specimen plots across groups. Compounds with significant between-group changes were determined based on variable effect on projection > 1 and P < 0.05 according to peak areas.

### Construction and validation of the predictive model for RAF

The LASSO method was employed for selecting the most efficient predictive indexes from distinctive taxa between the non-RAF and RAF groups [33, 34]. A taxonomic score (Tax-score) was determined for individual patients by linearly combining the retained taxa weighted by the corresponding coefficients. Predictive model performance was assessed using the AUC. Internal validation followed a reported protocol [37]. Meanwhile, the mean of 500 bootstrapped estimates of optimism was subtracted from the initial (full cohort model) estimate of the AUC and Nagelkerke R2 to obtain the bootstrap optimism-corrected estimates of performance. Lasso-logistic regression was carried out with the “glmnet” package in R. The “rms” package was utilized for multivariate binary logistic regression, nomogram construction and calibration graphs. C-index calculation was performed the “Hmisc” package. DCA was carried out with the “rmda” package. The optimal cut-off value for Tax score and Kaplan-Meier survival curves (with log-rank test) were obtained by the “survminer” and “survival” packages in R [70]. AUC, comparison, and NRI were obtained and calculated by the StataSE software.

### Statistical analysis

Normally distributed quantitative variables are mean±standard deviation (SD), and assessed by the t-test. Those with non-normal distribution were shown as median and quartiles, and compared by the Wilcoxon rank sum test. Qualitative data were shown as percentage, with the χ2 test for comparisons. Shannon index, Chao richness, and Pielou evenness were calculated with R version 3.3.3 in vegan package. PCA was carried out utilizing the FactoMineR package. PCoA used the vegan and ape packages. NMDS was carried out with the vegan package. Plot visualization utilized the ggplot2 package in R. Abundance differences at the gene, genus, species and KEGG module, respectively, were assessed by the Wilcoxon rank sum test, with Benjamini and Hochberg correction. Pearson correlation analysis was carried out for identifying microbiome-metabolome associations. Two-sided P < 0.05 indicated statistical significance.

## Data Availability

The data supporting the results of this article has been deposited in the EMBL European Nucleotide Archive (ENA) under the BioProject accession code PRJEB28384 [http://www.ebi.ac.uk/ena/data/view/PRJEB28384]. And the raw sequence data reported in this paper have been deposited in the Genome Sequence Archive (Genomics, Proteomics & Bioinformatics 2017) in BIG Data Center (Nucleic Acids Res 2019), Beijing Institute of Genomics (BIG), Chinese Academy of Sciences, under accession numbers CRA002277, that are publicly accessible at https://bigd.big.ac.cn/gsa. Metabolomics data can be found on the NIH’s Common Fund’s Data Repository and Coordinating Center website (Metabolomics Work-bench; Study IDs ST001168 and ST001169 for fecal and serum metabolomics, respectively).

## Competing interests

The authors declared no competing interests to this work.

## Author contributions

XCY, KBL, KZ, and JL carried out study conception, design, and supervision, as well as data interpretation and manuscript writing. JZ, PW, JJ, ZL, XDY and XQL were involved in patient enrolment, diagnosis, and data collection. KZ and JL performed data analysis. XCY, JZ and KBL carried out manuscript revision. The submitted manuscript had approval from all authors after reading.

## Acknowledgments

The present study was funded by the National Natural Science Foundation of China (81670214, 81500383, 81870308, and 81970271), the Beijing Natural Science Foundation (7172080), the Beijing Municipal Administration of Hospitals’ Youth Programme (QML20170303), and the 1351 personnel training plan (CYMY-2017-03).

## Ethics

The study had approval from the ethics committee of Beijing Chaoyang Hospital and Kailuan General Hospital and the signed informed consent was provided by each participant.

## Supplementary material

**Additional files 1: Fig. S1, Effects of baseline features (age, T2DM, total cholesterol, and drug utilization) on GM shift**

a. PCA evaluating age and microbial abundance at the genus level. A total of 90 specimens were assigned to 3 groups based in age (< 55 years, yellow; 55-65, light pink; > 65, violet). Squares represent non-AF CTR, triangles mean non-RAF and circles denote RAF.
b. PCA evaluating type 2 diabetes mellitus and microbial abundance at the genus level. A total of 90 specimens were assigned to 2 groups based on previous T2DM (no T2DM, grey; T2DM, dark purple). Squares represent non-AF CTR, triangles mean non-RAF and circles denote RAF.

a. PCA assessing TC and microbial abundance at the genus level. A total of 90 specimens were assigned to 2 groups based on TC levels (TC < 5.18, green; TC ≥ 5.18, purple). Squares represent non-AF CTR, triangles mean non-RAF and circles denote RAF.
b. PCA assessing drug and microbial abundance at the genus level. A total of 40 AF specimens were assigned to 11 groups based on used drugs: ACEIs, yellow triangles; ARBs, inverted green triangles; amiodarone, blue rhombi; statins, green asterisks; DMBG, red circles; an ACEI or ARB with amiodarone, brown hexagons; ACEI and statins, red hexagons; amiodarone and DMBG, dark red hexagons; ARB, amiodarone and statin, dark brown circles with forks; no medication, grey squares.

**Additional files 2: Table S1, Relative abundance levels of annotated genera**

**Additional files 3: Table S2, Relative abundance levels of annotated species**

**Additional files 4: Fig. S2, Altered taxonomic profiles from non-RAF to RAF**

a. Venn diagram depicting the count of annotated genera common to non-AF CTR (blue), non-RAF (pink) and RAF (red) cases. The overlap reveals 1219 genera simultaneously detected in non-AF CTR and AF with or without recurrence.
b. Venn diagram depicting the count of annotated species common in non-AF CTR (blue), non-RAF (pink) and RAF (red) cases. The overlap reveals 5041 species simultaneously detected in non-AF CTR and AF with or without recurrence.
c. Bar plots of relative abundance levels of the first 10 genera detected in the non-AF CTR (blue), non-RAF (pink) and RAF (red) groups. Distinct genera are depicted by different colors.
d. Bar plots of relative abundance levels of the first 10 species detected in the non-AF CTR (blue), non-RAF (pink) and RAF (red) groups. Distinct species are depicted by different colors.
e. Bar plots of relative abundance levels of the first 10 genera detected in non-AF CTR (blue), non-RAF (pink) and RAF (red) cases. Distinct genera are depicted by different colors.
f. Bar plots of relative abundance levels of the first 10 species detected in non-AF CTR (blue), non-RAF (pink) and RAF (red) cases. Distinct genera are depicted by different colors.

**Additional files 5: Table S3, Differential genera and species**

**Additional files 6: Table S4, Differential KEGG modules**

**Additional files 7: Fig. S3, Serum and fecal metabolic patterns in the control, non-RAF and RAF groups**

a, b. PLS-DA scores according to serum metabolic profiles in the non-RAF and RAF groups in ESI+ (a) and ESI− (b) modes. Pink triangles represent non-RAF and red circles denote RAF. A clear separation between non-RAF and RAF patients was obtained under both the ESI+ and ESI-modes.

c, d. OPLS-SA scores according to serum metabolic profiles in the non-RAF and RAF groups in the ESI+ (c) and ESI− (d) modes. Pink triangles and red circles represent non-RAF and RAF, respectively. The non-RAF and RAF groups were overtly separated under both the ESI+ and ESI-modes.

e, f. PLS-DA scores according to fecal metabolic profiles in the non-RAF and RAF groups in the ESI+ (e) and ESI− (f) modes. Pink triangles and red circles represent

non-RAF and RAF, respectively. The non-RAF and RAF groups were overtly separated under both the ESI+ and ESI-modes.

g, h. Orthogonal PLS-DA (OPLS-SA) scores according to fecal metabolic profiles in the non-RAF and RAF groups in the ESI+ (e) and ESI− (f) modes. Pink triangles and red circles represent non-RAF and RAF, respectively. The non-RAF and RAF groups were overtly separated under both the ESI+ and ESI-modes.

**Additional files 8: Table S5, Data of 8 metabolites with differential enrichment across groups**

**Additional files 9: Table S6, Taxonomic scores and CAAP-AF-Tax scores from non-RAF to RAF**

**Additional files 10: Table S7, Univariate and multivariable Cox regression analyses of factors potentially predicting AF recurrence**

